# Visualization of class A GPCR oligomerization by image-based fluorescence fluctuation spectroscopy

**DOI:** 10.1101/240903

**Authors:** Ali Isbilir, Jan Möller, Andreas Bock, Ulrike Zabel, Paolo Annibale, Martin J. Lohse

## Abstract

G protein-coupled receptors (GPCRs) represent the largest class of cell surface receptors conveying extracellular information into intracellular signals. Many GPCRs have been shown to be able to oligomerize and it is firmly established that Class C GPCRs (e.g. metabotropic glutamate receptors) function as obligate dimers. However, the oligomerization capability of the larger Class A GPCRs (e.g. comprising the β-adrenergic receptors (β-ARs)) is still, despite decades of research, highly debated.

Here we assess the oligomerization behavior of three prototypical Class A GPCRs, the β_1_-ARs, β_2_-ARs, and muscarinic M_2_Rs in single, intact cells. We combine two image correlation spectroscopy methods based on molecular brightness, i.e. the analysis of fluorescence fluctuations over space and over time, and thereby provide an assay able to robustly and precisely quantify the degree of oligomerization of GPCRs. In addition, we provide a comparison between two labelling strategies, namely C-terminally-attached fluorescent proteins and N-terminally-attached SNAP-tags, in order to rule out effects arising from potential fluorescent protein-driven oligomerization. The degree of GPCR oligomerization is expressed with respect to a set of previously reported as well as newly established monomeric or dimeric control constructs. Our data reveal that all three prototypical GPRCs studied display, under unstimulated conditions, a prevalently monomeric fingerprint. Only the β_2_-AR shows a slight degree of oligomerization.

From a methodological point of view, our study suggests three key aspects. First, the combination of two image correlation spectroscopy methods allows addressing cells transiently expressing high concentrations of membrane receptors, far from the single molecule regime, at a density where the kinetic equilibrium should favor dimers and higher-order oligomers. Second, our methodological approach, allows to selectively target cell membrane regions devoid of artificial oligomerization hot-spots (such as vesicles). Third, our data suggest that the β_1_-AR appears to be a superior monomeric control than the widely used membrane protein CD86.

Taken together, we suggest that our combined image correlation spectroscopy method is a powerful approach to assess the oligomerization behavior of GPCRs in intact cells at high expression levels.

## Introduction

G protein-coupled receptors (GPCRs) constitute the largest class of membrane-bound receptors with >800 members expressed in humans. GPCRs relay extracellular stimuli into intracellular signals and modulate almost every physiological process. The fidelity of GPCR signaling is fine-tuned on three different levels. First, GPCRs can possess distinct binding sites for extracellular ligands and some receptors get activated by multiple endogenous ligands. Second, intracellular adaptor proteins such as G proteins, GRKs and β-arrestins may channel the extracellular signal into distinct cellular outcomes. Third, GPCRs signaling can be modulated within the plasma membrane by forming dimers and/or higher-order oligomers [1–5].

Whereas Class C GPCRs function as obligate dimers [6], the situation for the larger family of Class A GPCRs is less clear. Although Class A GPCRs are fully functional as monomers [7], there are multiple lines of evidence that Class A GPCRs can also form dimers[4]. However, the impact of dimerization on GPCR function and signaling, is not completely understood. As receptor monomers and dimers may have distinct functions on cell signaling, it is important to rigorously assess the dimerization behavior in intact cells with appropriate methods.

Since the first bioluminescence resonance energy transfer (BRET) investigation on the oligomerization behavior of β_2_-adrenergic receptors (β_2_-ARs) [8], a large number of studies have addressed the oligomerization state of GPCRs with fluorescence approaches [4].

However, there are conflicting data on the existence and abundance of GPCR dimers in intact cells. In the extreme case, by using the same method and the same receptor, such as in the case of BRET read-out of β_2_-AR oligomerization, certain reports support the presence of oligomers [8], while others suggest that they are absent [9].

One of the most promising and accurate approach developed over the last few years appears to be single molecule tracking (SMT) [10]. SMT provides precise information on single molecule dynamics while at the same time offering insight into the specific oligomerization state of a protein by using the intensity of the localized spots. The major limitation of the method is the assessment of oligomerization at plasma membrane concentrations exceeding a few receptor molecules/μm^2^ [11, 12]. At concentrations above this value, individual molecular point spread functions (PSF) begin to significantly overlap and accurate localization and tracking becomes impossible. This limits the exploration of higher expression levels, which, although not always physiological, provide an interesting range where to test the law of mass action and compare experimental data to predictions arising from coarse-grained molecular simulations, which tend to be performed at much higher “in-silico” concentrations [13]. Moreover, the majority of reports on GPCR dimerization have been performed in overexpressed systems.

Fluorescence fluctuation spectroscopy techniques offer an effective tool to investigate with good precision receptor dimerization at higher expression levels than SMT. Single point fluorescence correlation spectroscopy was used in the past to characterize GPCR diffusion, formation of hetero-complexes [14] and receptor homo-dimerization, by analyzing the histogram of the collected photons [15, 16]. Time-based image fluorescence fluctuation spectroscopy methods, which rely on the statistical analysis of many pixels of an image, have allowed to characterize the oligomerization state of the GPI-anchored membrane receptor uPar [17] and the ErbB [18] observing the agonist-dependent formation of dimers and oligomers. More recently, GPCRs such as the 5-HT_2C_ and the muscarinic M_1_ receptor [19, 20] were investigated by spatial intensity distribution analysis (SpIDA): the authors observed an antagonist-promoted oligomerization in the case of M_1_ receptors and, in contrast, an antagonist-dependent disruption of 5-HT_2C_ receptor oligomers.

The fundamental advantage of spatial-temporal brightness analysis over SMT is that a spatially resolved view of the plasma membrane allows discarding from the analysis those regions where receptor aggregation phenomena different than oligomerization, such as recruitment by ‘endocytic machinery’, have occurred [21, 22].

In this report, we combine two image-based fluctuation spectroscopy methods, namely temporal brightness (TB) [17, 23, 24] and SpIDA [25] to characterize the oligomerization state of three prototypical GPCRs, the homologous β_1_-AR and β_2_-ARs, and the muscarinic M_2_ receptor (M_2_R) at expression levels of the order of tens to hundreds of receptors/μm^2^, a concentration level where oligomerization driven exclusively by physical kinetics should have already occurred. By comparing the receptors with a set of monomeric and dimeric reference proteins, and by employing two labeling strategies based on C-terminal fluorescent protein fusions and N-terminal SNAP-tag labels, we find here clear evidence of a predominantly monomeric state for all these three receptors in intact cells. The absence of a dominant dimeric fraction at these concentrations suggests that kinetic oligomerization is inefficient, characterized by slow on-rates and/or fast off-rates, resulting in a short lifetime of any oligomeric complex.

## Results

### C-terminally EYFP-labeled controls

In order to assess the oligomeric nature of β_1_-ARs, β_2_-ARs and M_2_Rs we first devised a set of reference proteins to characterize the brightness of constitutively monomeric and dimeric EYFP-tagged membrane proteins [17]. Such controls carry the same fluorophore as the target constructs, and should share a similar diffusion coefficient and mode of motion (Supplementary Figure 1).

**Figure 1:**
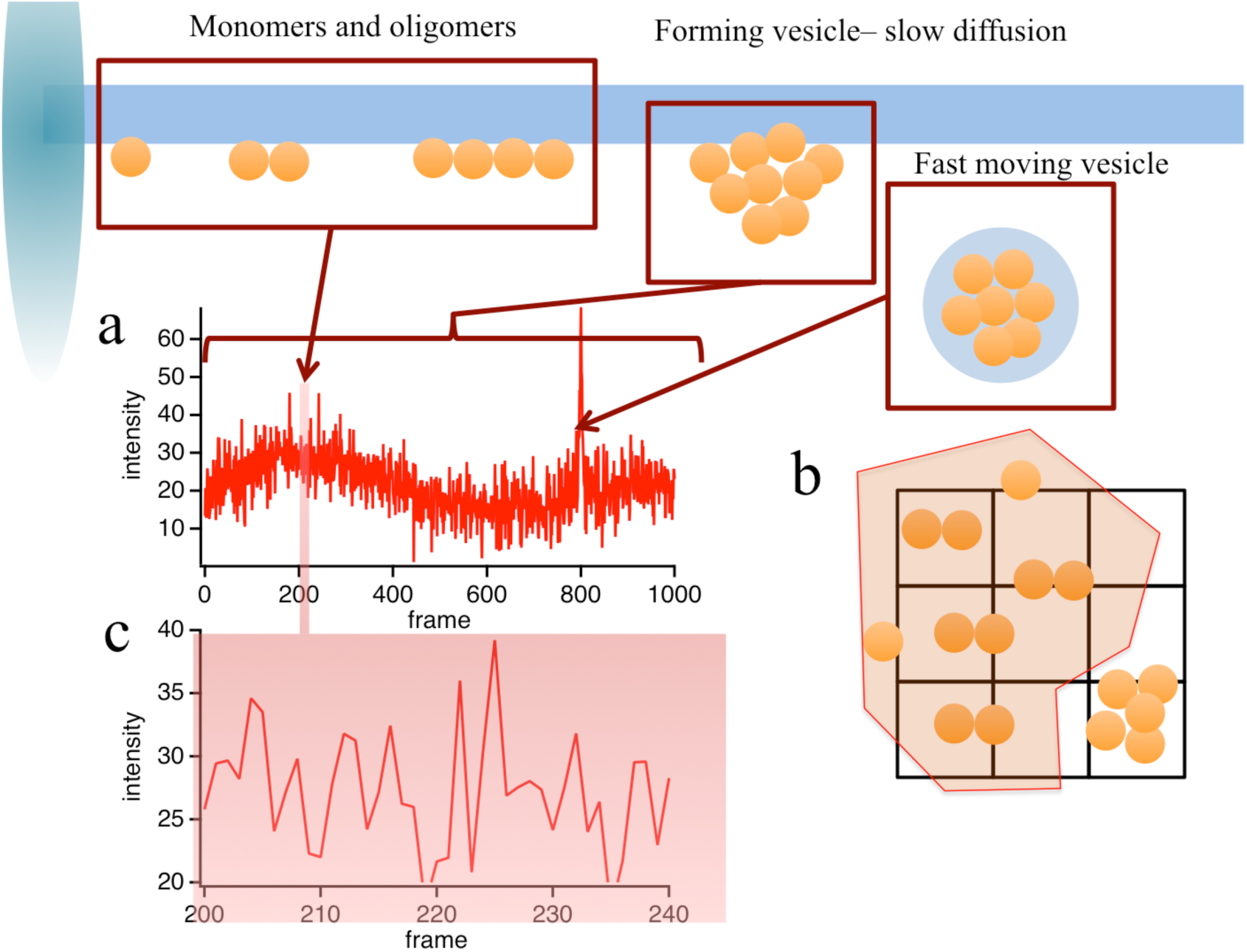
Graphical representation of the impact of different oligomerization and other dynamic states of the labeled receptors on the measured molecular brightness. a) The measured intensity values measured on a pixel as a function of time or b) across many pixels reflect also the presence of higher order aggregates and the dynamics of processes occurring on a completely different timescale compared to molecular diffusion. c) For this reason, fluorescence fluctuations arising from molecular diffusion have to be extracted in time or in space by excluding spurious processes. The combination of the two approaches, spatial and temporal brightness proves here a powerful tool to extract molecular brightness values reflecting the true oligomerization state the membrane receptors.

Based on previous reports we identified the single transmembrane peptide CD86 (also known as B7–2) as a potential monomeric control. This construct is routinely used as a monomeric reference in photo activated localization microscopy (PALM) experiments [26] and was previously used by our group to calibrate SMT experiments [12]. We chose to tag all our constructs C-terminally with EYFP (ex. 513 nm, em. 527 nm, EC 83400, QY 0.61) as we found no evidence for intrinsic dimerization of EYFP in comparison to monomeric YFP (mYFP) [27] (Supplementary Figure 2).

**Figure 2.**
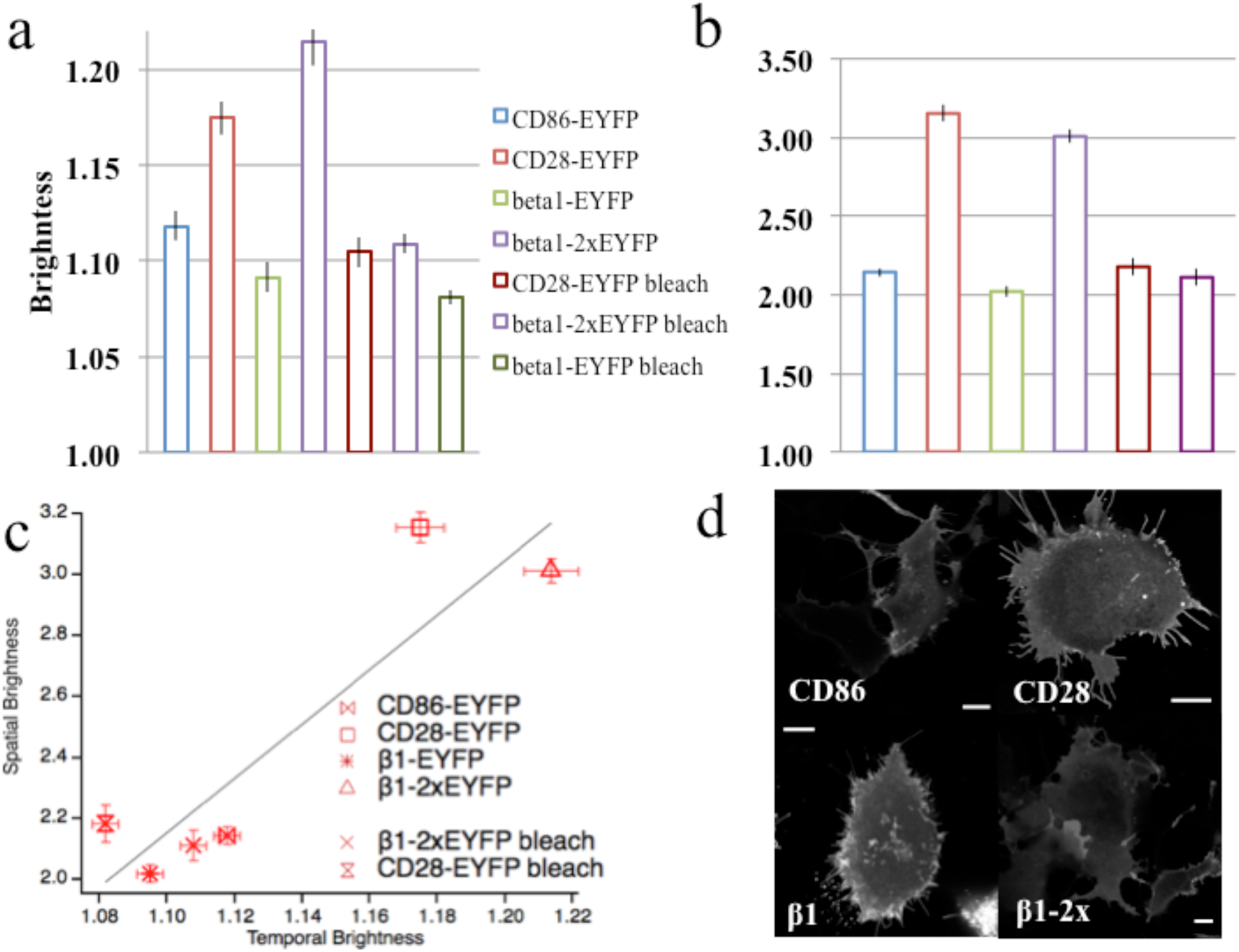
Brightness Analysis of C-terminally EYFP-tagged constructs. a) Temporal Brightness measured on a set of control constructs. CD86-EYFP is used as the monomeric reference. CD28-EYFP is used as the dimeric reference. CD86-EYFP and β_1_-AR-EYFP were both tested as monomeric controls. β_1_-AR-2xEYFP serves as the reference of a constitutive dimer. Decrease in brightness in photobleached samples of β_1_-AR-2xEYFP and CD28-EYFP reveal the oligomeric nature of the controls. Image sequences (50–100 frames) were acquired at a line rate of 400 Hz, 256×256 pixels. Pixel size = 50 nm b) Corresponding spatial brightness measurements on the same set of controls. The image was acquired at 100 Hz, 512×512 pixels. Pixel size = 50 nm. c) Correlation between SpIDA and Temporal Brightness measurements. The measured slope is 8.9, which accounts for the higher apparent brightness measured in SpIDA experiments: 12x higher laser power, 4 times longer dwell time. d) Images of HEK293-AD cells transfected with C-terminally labeled EYFP constructs, showing correct basal membrane localization of the constructs and a relatively homogeneous membrane distribution. Scale bars are 10 μm.

Another important characteristic that the control construct should exhibit is a homogeneous expression on the plasma membrane. If this is not the case, and a large number of aggregates such as forming vesicles and mature endosomes are present, then both spatial and temporal brightness measurements may be affected (Supplementary Figure 3). In our hands, using HEK293-AD cells, CD86 displayed a robust cell membrane expression but was also, albeit to a lesser extent, found in cytosolic and near-membrane aggregates dotting the basal membrane (**Figure 1**). The presence of such aggregates may affect the correct brightness readout (e.g. an endosome will appear –in brightness-as a large oligomer (Figure 1a), and moving vesicles can generate extra variance over space and time). This is one of the reasons why we decided to employ two complementary approaches to measure molecular brightness: measurement of brightness over time (TB) and over space (SpIDA). Large, immobile or slow (nm/s) and bright features can be easily treated in TB analysis by a boxcar filter detrend [28], while they have to be avoided from the region of interest (ROI) used to extract SpIDA values (Figure 1b). On the other hand, since SpIDA brightness values can be extracted from one image, this latter approach is less sensitive to photobleaching, drift or defocus of the sample. Considering the different acquisition parameters, in particular, the pixel dwell time, spatial and temporal brightness values are expected to be distinct. The average apparent brightness of CD86-EYFP measured on a stack of 50–100 images (from the variance of the intensity of each pixel over time, i.e. temporal brightness) and on a pool of 21 cells is 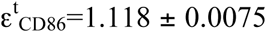 (Figure 2a). The SpIDA analysis of CD86 yielded a brightness value of 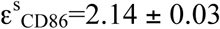 (Figure 2b).

**Figure 3.**
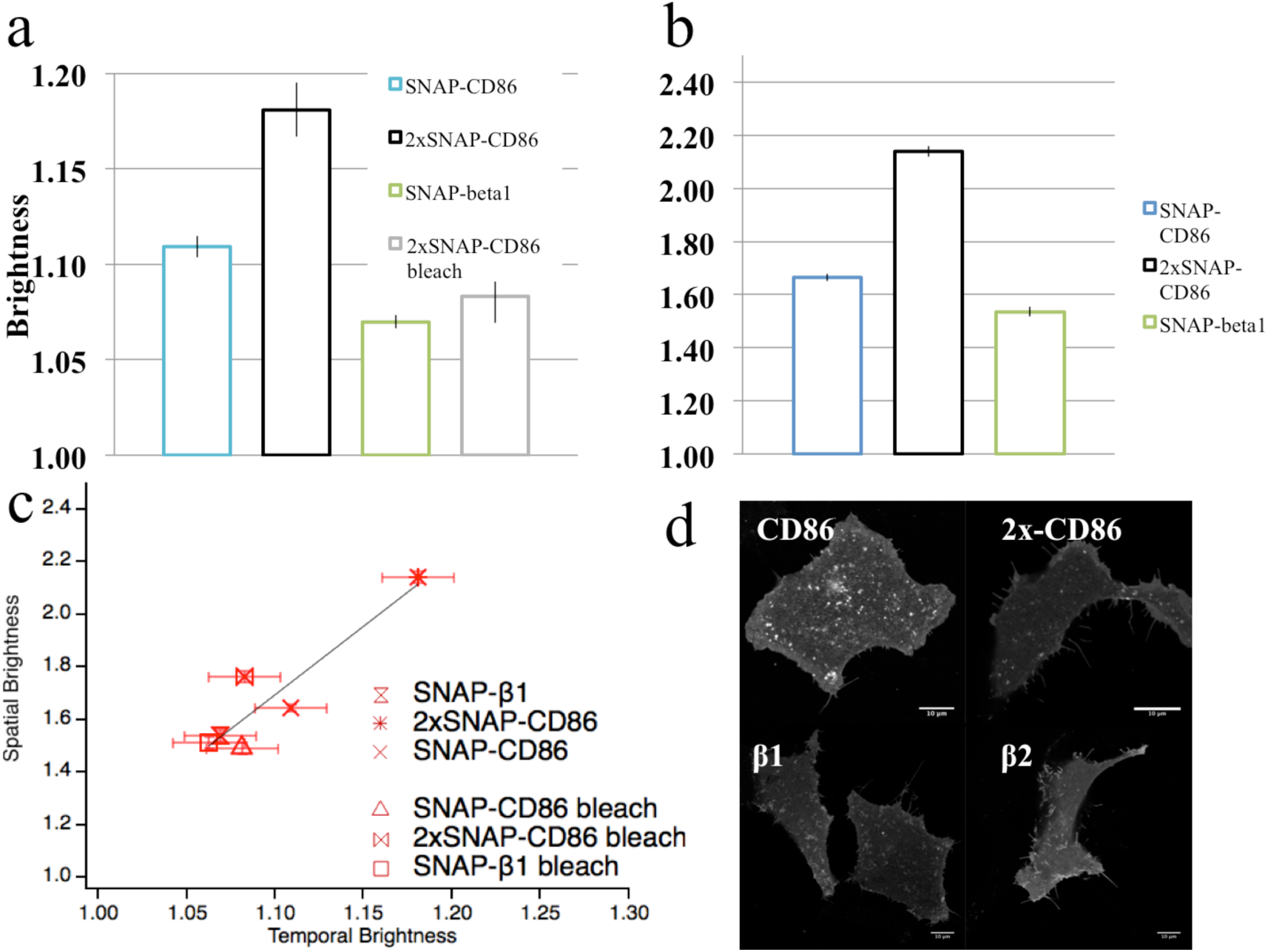
Brightness Analysis of N-terminally SNAP-tagged constructs, labeled with SNAP-surface 488. a) Temporal Brightness measured on the set of control constructs. SNAP-CD86 is tested as a monomer reference. 2xSNAP-CD86 is used as a dimer reference. SNAP-β_1_-AR displays a brightness lower than that of SNAP-CD86. Decrease in brightness in photobleached samples of 2xSNAP-CD86 reveals the oligomeric nature of the control. b) Corresponding spatial brightness measurements on the same set of controls. c) Correlation between SpIDA and Temporal Brightness measurements. The measured slope is about 5, which partially accounts for the higher apparent brightness measured in SpIDA experiments: 12x higher laser power, 4 times longer dwell time. d) Images of HEK293-AD cells transfected with the appropriate SNAP-tagged constructs, showing correct basal membrane localization of the constructs and a relatively homogeneous membrane distribution. Scale bars are 10 μm.

To obtain a reference brightness dimer, we pursued two strategies. In the first approach, we relied on CD28, a transmembrane protein displaying a disulfide-linked dimeric structure. The temporal and spatial brightness obtained for CD28 are 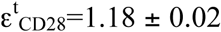 and 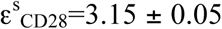, respectively (Figure 2 a, b). Both temporal and spatial brightness values are smaller than twice the values for 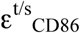, viz. 1.24 and 3.28.

We then checked a reference GPCR, which in previous investigations from our own group, [12, 29] displayed a monomeric fingerprint, the β_1_-AR. Interestingly, β_1_-AR-EYFP displayed an apparent brightness lower than that of CD86-EYFP, 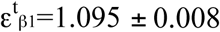 and 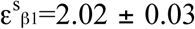 (Figure 2a, b). These values are compatible with half the brightness for CD28-EYFP, suggesting that CD86 may dimerize to a certain extent in intact cells. This would be in line with previous findings of a preferred rather than exclusive monomeric organization of CD86 [30].

The second approach to generate a dimeric control was to C-terminally tag GPCRs with two EYFP molecules separated by a single α-helical spacer to limit self-association of EYFP molecules (see Materials and Methods and Supplementary Figure 4e). The brightness measured for this β_1_-AR-2xEYFP construct is 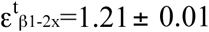 and 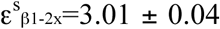, in full agreement with the values measured for CD28 and twice the values measured for β_1_-AR-EYFP.

**Figure 4.**
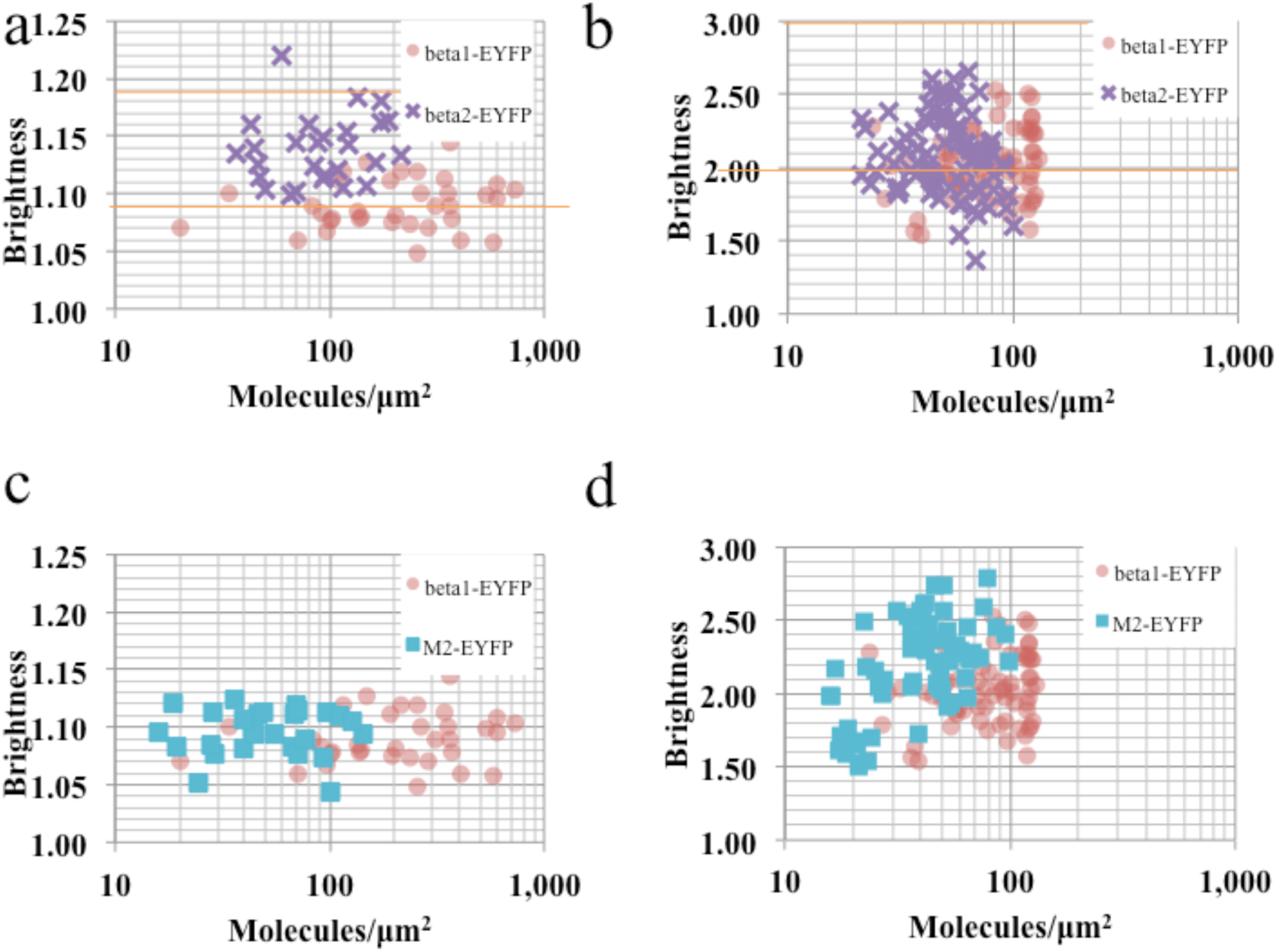
Brightness-expression plots for three prototypical GPCRs: β_1_-AR, β_2_-AR, and M_2_R, either C-terminally EYFP or N-terminally SNAP-tagged. a) Temporal brightness analysis of β_1_-AR-EYFP (•) and β_2_-AR-EYFP (x). β_1_-AR-EYFP brightness defines the brighntess of a monomer (horizontal line at 1.10). β_2_-AR-EYFP cells display a higher average brightness, but far from that of a constitutive dimer (horizontal line at 1.21). Each point represents a cell. b) Spatial brightness analysis of β_1_-AR-EYFP (•) and β_2_-AR-EYFP (x). β_1_-AR-EYFP brightness matches that of the monomer. Each point represents a ROI within a cell, 3 ROIs per cell. Horizontal lines are defined as in a c) Temporal brightness analysis of β_1_-AR-EYFP (•) and M_2_R-EYFP (■). M_2_R-EYFP displays a lower overall expression, but the brightness is also in agreement with that of a prevalent monomeric species. d) Spatial brightness analysis of β_1_-AR-EYFP (•) and M_2_R-EYFP (■). Each point represents a ROI within a cell, 3 ROIs per cell.

To further test the quality of our measurements, we subjected cells expressing the dimeric constructs CD28 and β_1_-AR-2xEYFP to whole cell photobleaching. Due to the stochastic nature of photobleaching, many of the fluorophores of a dimer will be photobleached (although the actual physical dimer is not affected), resulting in an apparent increase of the monomer/dimer ratio. After sustained photobleaching, we observe that the apparent brightness of CD28 and β_1_-AR-2xEYFP revert to the approximately monomeric value: 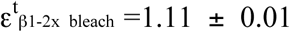 and 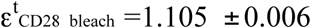, while for SpIDA we obtain 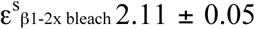 and 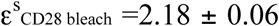. (Figure 2 a, b). In contrast, when bleached, β_1_-AR-EYFP displays a negligible reduction in brightness (Supplementary Figure 6a).

The agreement between temporal and spatial brightness measurements is overall excellent. In our experimental conditions the brightness measured by SpIDA is 9 times larger than what we measure by TB, which accounts for the higher laser power (8x) and lower scan rate (0.25x) used in spatial brightness measurements (Figure 2c).

Finally, all our controls displayed correct membrane localization, as illustrated in the panels of Figure 2d, as well as a diffusion coefficient in agreement with previous observations, in the range of 0.1 μm^2^/s (Supplementary Table 1).

### N-terminally SNAP-tagged controls

In order to minimize potential contributions from cytosolic aggregates containing EYFP, we decided to further validate our findings with an alternative labeling strategy. Towards this goal, we worked with N-terminally SNAP-tagged constructs labeled with cell membrane impermeable SNAP dyes. Using this approach, the extracellular SNAP-tag is labeled by incubating the cells with an organic dye, in our case Atto488 (See Materials and Methods). We first tested SNAP-CD86 and a double SNAP-tagged construct, 2xSNAP-CD86 as monomer and dimer controls, respectively. We recorded 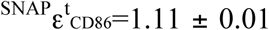 and 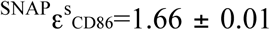 (Figure 3a, b). The dimer reference 2xSNAP-CD86 yielded 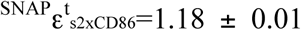 and 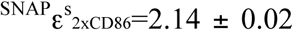. We observe again that the dimer falls short of double the brightness for SNAP-CD86 in both types of measurement. However, when we measure the brightness for SNAP-β_1_-AR we observe a lower value than for SNAP-CD86: 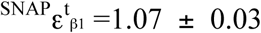 and 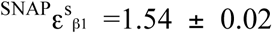. This value is now fully compatible with half the values measured for our dimeric control. After sustained photobleaching, we observe that the apparent brightness of 2xSNAP-CD86 reverts to the approximately monomeric value: 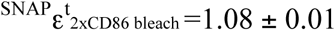.

The SNAP-tagged constructs displayed a similar cellular localization as their C-terminally-tagged EYFP counterparts and we find an excellent photostability, as displayed in Supplementary Figure 4b.

### Oligomerization state of three prototypical GPCRs

Within our controls, the β_1_-AR displays a pronounced monomeric character, since it is characterized by a molecular brightness lower than CD86, and this is independent of the type of labeling (C-terminal EYFP or N-terminal SNAP-tag). Photobleaching of EYFP-labeled β_1_-AR yielded post-bleach values of 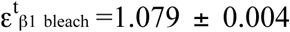 and 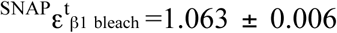 for SNAP-tagged receptors, which are both compatible with the pre-bleach values, suggesting that oligomerization is negligible.

Given the fact that brightness values are a weighted average of the brightness of each species present in a pixel, immobile background fluorescence skews any ε value towards one. For this reason, we considered in our averages only cells displaying a cell membrane expression in excess of 10 molecules/μm^2^. However, the same conclusions concerning the bleaching experiments apply to β_1_-AR also for expression levels below this threshold. Full brightness *vs.* expression data are displayed for β_1_-ARs in Figure 4.

We used β_1_-ARs as a reference to compare the constitutive brightness *vs.* expression curves of the β_2_-AR and the M_2_R. Our findings indicate that β_2_-ARs form more dimers than β_1_-ARs, in agreement with our previous findings [12]. The average temporal and spatial brightness of 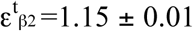 and 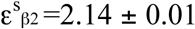, respectively, indicate a mixture of monomers, dimers, and, potenitally higher order oligomers, but entirely exclude the possibility that β_2_-ARs form constitutive dimers (Figure 4 a,b).

M_2_Rs displayed also a largely monomeric fingerprint when observed by temporal brightness, as the brightness values of M_2_R-EYFP largely overlap with those measured for β_1_-AR-EYFP. The average brightness value is 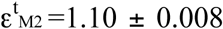. On the other hand, spatial brightness reveals a small fraction of dimers for this receptor, since 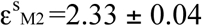 and post-bleach we observe 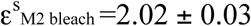.

As a control, we investigated SNAP-β_1_-AR and SNAP-β_2_-AR. The temporal brightness measurement confirmed the higher oligomerization state of SNAP-β_2_-ARs, as displayed in Supplementary Figure 4 c. We observe 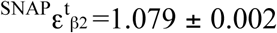, which compares to 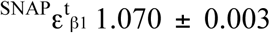 of SNAP-β_1_ (Supplementary Figure 5a). Upon photobleaching, the brightness of SNAP-β_1_-AR was virtually unchanged (Supplementary Figure 6b). We are, however, far from the constitutive dimer value corresponding to 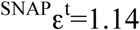. SpIDA displayed comparable results, yielding 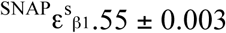 and 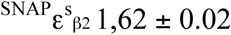 Supplementary Figure 5b.

## Discussion

We report here an experimental protocol based on two different methods and two labeling strategies to address the longstanding controversy of GPCR oligomerization in intact cells when high expression levels of the receptors are observed. Our experimental strategy, based on fluorescence fluctuation spectroscopy methods, allows measuring the oligomerization state of fluorescently labeled receptors in living cells. The first experimental approach, temporal brightness analysis, is an image based version of the Photon Counting Histogram method [23], which was recently used to characterize the oligomerization state of many Class A GPCRs, including the β_2_-AR and M_2_R used in our study [15]. The second approach, SpIDA, based on the statistical analysis of the pixel intensities in a single LSM image, has been extensively used recently to assess GPCR oligomerization, providing indication for the oligomerization state of the 5-HT_2C_ and the M_1_R [19, 20].

The results from both methods are in an overall excellent agreement (Figure 2c, Figure 3c). The comparison of two alternative labeling strategies (intracellular fluorescent proteins and extracellular SNAP-tags) confirmed our findings, ruling out the possibility that oligomerization may be driven by the fluorescent tag itself. Furthermore, given the rather intricate and heterogeneous morphology of the basal membrane of adherent cells, the use of an image-based method allowed us discriminating regions of the membrane where a homogeneous expression of the receptor was present from those were it was not, and large aggregates or other dynamic processes than receptor diffusion were generating brightness gradients or hot-spots (**Figure 1**a and Supplementary Figures 3 and 4). This effect can be clearly seen by observing how the measured brightness changes upon a modification of the area of analysis. By progressively enlarging the area of interest, the measured brightness increases as endosomes and other bright features of the plasma membrane are included in the intensity histogram (Supplementary Figure 7).

While the photon counting histogram is a powerful method which allows discriminating the number and brightness of up to two existing species mixed within a homogeneous sample [23], it is insensitive to such heterogeneities: if not properly corrected, the histogram of the photon counts may be affected by fluctuations that do not originate from the receptor diffusion within the membrane (**Figure 1**a,b). As far as GPCRs are concerned, another important phenomenon that may affect the apparent oligomerization is receptor internalization. The initial step of the internalization process is the segregation of multiple receptors within coated pits. Clathrin coated pits are hubs were receptors can cluster, which display a rather slow diffusion coefficient, at least an order of magnitude slower than the receptor. While mature endosomes in the sub-membrane region are large and bright enough to be identified and accounted for in the analysis (**Figure 1**c), small, forming pits may render the data intractable for a proper brightness analysis. According to this reasoning we refrained here from studying GPCRs under any kind of pharmacological stimulus besides the use of the inverse agonist ICI 118551 to counteract internalization and enhance the amount of β_2_-ARs expressed on the plasma membrane.

While internalization is present also in basal conditions, and may vary according to the receptor under study [22], this did not skew significantly our results: the controls using N-terminally SNAP-tagged constructs, imaged immediately after labeling, provided results consistent with what we observed using C-terminal EYFP. This observation, together with the finding that the brightness of a construct carrying EYFP was comparable to that of the same construct labeled with mYFP, the monomeric version of EYFP (Supplementary Figure 2), confirmed the validity of our approach.

The SNAP-tag labels displayed an overall excellent photostability (negligible photobleaching over the 50–100 frames used for imaging, under our experimental conditions), making them ideal choices for temporal brightness measurements where photobleaching may be a problem. Interestingly, we obtained similar or even lower brightness values when using Atto488-labeled SNAP-tagged constructs compared to EYFP. Nominally Atto488 has a 1.5x higher brightness than EYFP, and the laser power at 488 nm employed to excite it was 2.5x greater than the corresponding laser power at 514 nm used to excite EYFP. The reason that the expected gain in brightness was not observed is because the benzylguanine moiety conjugated to the Atto488 acts as a potent quencher, reducing the apparent brightness of the dye of a factor of seven [31]. This is an interesting, often overlooked feature of SNAP-dyes, since, depending on the molecular structure of the dye, strong quenching may occur after benzylguanine conjugation.

In the case of M_2_R-EYFP, SpIDA measurements reported a higher 8 value than TP measurements. We speculate that the organization of the basal membrane of M2R expressing cells is characterized by the presence of many, sub-diffraction limit vesicles (such as coated pits). Therefore it is not possible to exclude them from the area selection in SpIDA, whereas their effect on brightness can be filtered out in the TP measurements. Another explanation is that the observed TB values are lowered by the presence of a fraction of immobile M_2_Rs, possibly suggesting M_2_R-specific internalization dynamics.

Overall, our measurements fall within a rather heterogeneous background of previous measurements of GPCR oligomerization. In particular, for the three receptors that we investigated, previous research showed either a largely monomeric [32],[29], constitutive dimeric [15] or higher order oligomeric state [29, 33, 34]. In one case, concentration-dependent effects were observed for both β_1_-ARs and β_2_-ARs, with a different equilibrium constant depending on the receptor [12].

How are our data comparable with the previous results? Our measurements are in favor of a predominantly monomeric organization for these three receptors. As far as the β_1_-ARs are concerned, this observation is in agreement with most of the previous results [12, 29]. The only reports supporting a higher oligomerization state for β_1_-ARs come from whole-cell BRET measurements [33] and the results may have been influenced by receptor interactions not on the plasma membrane.

For M_2_Rs, single molecule imaging data [32] suggest a predominantly monomeric fingerprint, whereas fluorescence lifetime-based FRET measurements have indicated tetramers [34]. Photon counting histogram analysis indicates constitutive dimers [15]. Our observation of monomers is in line with another study in cardiac muscle [32]. As far as β_2_-ARs are concerned, our data show a higher degree of oligomerization, but exclude the presence of constitutive dimers. We worked at expression levels ranging from about 0.1 to 10 nM, corresponding to a surface expression level on the plasma membrane ranging from tens to hundreds of molecules/μm^2^. While these numbers are much larger than those used in SMT experiments [11, 12], they are in the range of previous reports using FCS and FRET/BRET techniques [8, 35].

In conclusion, we have analyzed the basal oligomerization state of three prototypical GPCRs, observing a largely monomeric state for two of them, M_2_Rs and β_1_-ARs, and a mixture of monomers and oligomers for β_2_-ARs. Our results point to the advantage of using image-based fluorescence fluctuation methods to assess the oligomerization state of class A GPCRs, and provide a way to discriminate between effects arising from receptor diffusion and higher order aggregates in the sub-membrane region.

## Materials and Methods

### Molecular Cloning

Plasmids coding for CD86-EYFP, CD28-EYFP, β_1_-AR-EYFP, β_2_-AR-EYFP were previously described [29].

In order to generate the β_1_-AR-2xEYFP construct, the β_1_-AR cDNA was amplified by PCR using the forward 5’-AATAATAAGCTTATGGGCGCGGGGGTGCTC-3’ and reverse 5’-AATAATGGATCCCACCTTGGATTCCGAGGCGAA-3’ primers and subcloned into pcDNA3.1. Two EYFP cDNAs, the second one with a stop codon, were sequentially subcloned to the C-terminus of the β_1_-AR after PCR amplification using primers for EYFP1:

forward 5’-AATAATGGATCCGTGAGCAAGGGCGAGGAG-3’ and reverse 5’-AATAATGAATTCCTTGTACAGCTCGTCCATGCC-3’ and for EYFP2: forward 5’-AATAATTCTAGAGTGAGCAAGGGCGAGGAGCTG-3’ and reverse 5’-AATAATGGGCCCTTACTTGTACAGCTCGTCCATGCC-3’.

A sequence encoding a single alpha helical linker (A(EAAAK)_4_A) was inserted between two EYFP sequences in order to prevent random interactions of two EYFPs within the same protein [36].

The mYFP construct was a kind gift of Roger Y. Tsien (University of California,San Diego, USA). C-terminally mYFP-tagged constructs were generated by deleting EYFP from respective constructs by restriction enzyme digestion using Xbal and Notl, and then subcloning the PCR-amplified mYFP sequence using the forward 5’-AAT AAT TCT AGAGT GAGCAAGGGCGAGGAGCTG-3’ and reverse 5’-AATAATGCGGCCGCTTACTTGTACAGCTCGTCCATGCC-3’ primers.

Plasmids coding for N-terminally SNAP-tagged CD86, CD28, β_1_AR, β_2_AR and 2xSNAP-CD86 were previously described [12].

### Cell Culture

All experiments were performed with transiently transfected HEK293-AD cells (Cell Biolabs, San Diego, USA). Cells were cultured in DMEM (Dulbecco’s modified Eagle’s medium) (PAN Biotech, Aidenbach, Germany), supplemented with 4,5 g/L Glucose, 2 mM L-Glutamine, 10% FCS (Biochrome), 100 units/mL penicillin and 0,1 mg/mL streptomycin and maintained at 37 °C and 5% CO_2_. Cells cultured in 15-cm dishes were split at a 1:36 ratio into 6-well plates containing poly-D-lysine (PDL)-coated 24 mm glass coverslips.

### Transient transfection

Cells were transfected using Effectene Transfection Reagent (QIAGEN, Hilden, Germany) according to the manufacturer’s instructions. Cells seeded on PDL-coated coverslips in 6-well plates were transfected 16 hours after seeding with 0.6 μg plasmid/well.

### SNAP-labeling

Cells transfected with SNAP-tagged receptors were labeled using the SNAP-Surface 488 Dye (New England Biolabs, Frankfurt am Main, Germany) according to the manufacturer’s instructions. Briefly, 24 hours after transfection, the culture medium was exchanged with labeling medium (5 μM SNAP-dye in DMEM) followed by a 30 minute incubation at 37°C. SNAP-Surface 488 was observed to localize in internal cytosolic compartments in untransfected cells, but not in the plasma membrane (Supplementary Figure 8).

### Spatial Intensity Distribution Analysis (SpIDA)

Transfected HEK293-AD cells grown on PDL-coated coverslips were placed into a custom designed cell chamber. Coverslips were washed once with HBSS (Hank’s Balanced Salt Solution) and the chamber was filled with 500 μL HBSS. The cell chamber was mounted onto a Leica SP5 confocal laser scanning microscope. Cells were imaged with a 40x / 1.25 numerical aperture oil immersion objective. A 65 mW Argon laser was set to 37% and a 488 nm laser line was used at 0.1% power in order to search for the cells expressing EYFP-tagged receptors. 512 × 512 pixel images of basolateral cell membranes expressing EYFP-tagged receptors were acquired with a Hybrid detector in photon counting mode, using a 514 nm laser at 10% power for excitation, and the respective emission was measured within 520–600 nm. The zoom factor was set to 15.15 x in order to reach a pixel size of 0.05 μm and the laser scanning speed was set to 100 Hz. For photobleaching experiments, a series of 3 images was acquired with the same acquisition settings. The third image was used for the analysis of brightness after photobleaching. A maximum of two cells was analyzed within each image.

Image analysis was preformed applying the SpIDA function (one-population mode) using a MATLAB routine, described previously [25]. The routine is courtesy of Antoine Godin (CERVO, Brain Research Centre - Laval University, Canada). Pre-bleaching and post-bleaching images were separately executed using FIJI. The ROIs for image analysis using SpIDA were drawn carefully in free area selection mode, implemented to the original SpIDA function, in order to avoid vesicles and inhomogenously distributed membrane areas more effectively [19]. Free area selection is superior to square or rectangular ROI selections because it allows the user to select homogenously distributed fluorescent particles even within an image that contains high content of vesicles and/or inhomogenous basolateral membrane areas. SpIDA analysis was performed on ROIs of four different area sizes ranging from ~10 to ~400 μm^2^. From each image, three ROIs were analyzed. For each experimental group, the number of analyzed cells ranged between 24 and 43. Number of molecules and brightness value were used from each ROI analysis to calculate fluorescence intensity and molecular concentration. The molecular concentration of EYFP-tagged receptors was calculated by dividing the mol (calculated by multiplying the number of molecules with the Avogadro number) by the focal volume.

### Temporal Brightness

For TP imaging the same setup was used as for SpIDA measurements. The imaging mode was XYT and 50 frames were taken with a scanner speed of 400 Hz using the following parameters: pinhole-size: 67.93/ zoom-factor: 30.3x/resolution 256×256 pixels. SNAP-labeled constructs were imaged using a 488nm laser power of 5% and the Hybrid detector was set between 520 and 600 nm. To perform photobleaching experiments the laser power between two imaging stacks was increased to 15% during 10 frames. EYFP-tagged constructs were imaged using a 514 nm laser power of 2.5 % and photobleached with 10 % laser power over 10 frames. Data were analyzed using a custom-written Igor Pro routine as described previously [28]. The brightness values were calculated based on the average of the brightness values from each pixel within the region of interest. This approach is equivalent to measuring the peak of the brightness histogram, as illustrated in Supplementary Figure 9.

## Acknowledgments

We are grateful to Antoine Godin (CERVO, Brain Research Centre - Laval University, Canada) for discussion concerning SpIDA data analysis. We would like to acknowledge the contribution of the students of the Master in Biohysics programme of the University of Würzburg, Germany, as well as that Jana Wächter and Sofia Krohne for the work performed during their internships.

## Contributions

*Performed Experiments*: Ali Isbilir, Jan Moeller, Paolo Annibale; *Analyzed Data:* Paolo Annibale, Ali Isbilir; *Wrote the Manuscript:* Paolo Annibale, Andreas Bock; *Contributed Reagents and Materials:* Ali Isbilir, Paolo Annibale, Ulrike Zabel, Jan Moeller; *Conceived and planned the experiments:* Paolo Annibale, Martin J. Lohse, Ali Isbilir, Jan Moeller, Andreas Bock; *Initiated the project:* Martin J. Lohse

---

**SI Figure 1.**
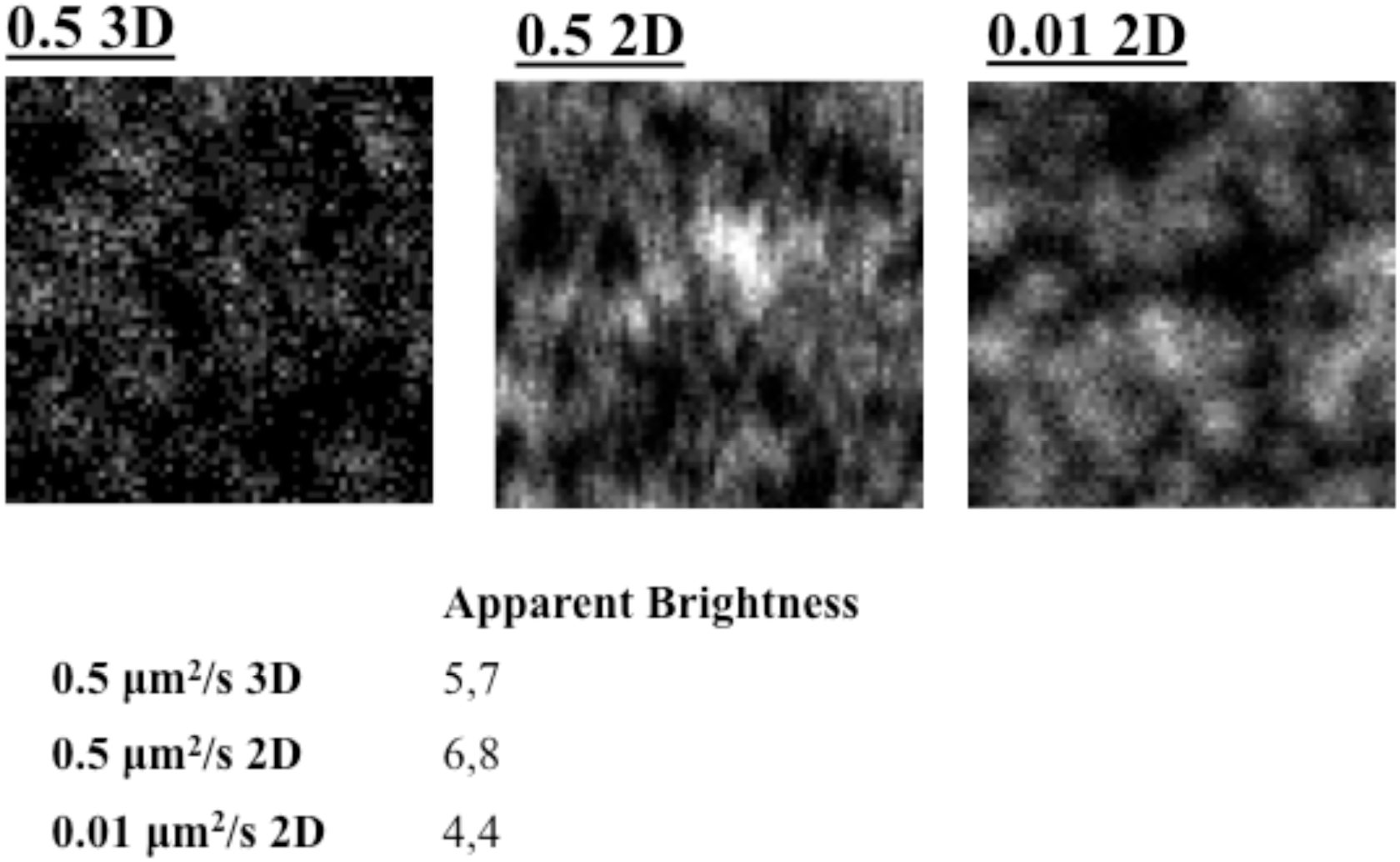
Simulated datasets containing fluorescent particles moving with different diffusion coefficients, either in a 3D or in a 2D space. The apparent brightness is not only function of the diffusion coefficient, but also, sizably, of the dimensionality of the space where diffusion takes place.

**SI Figure 2.**
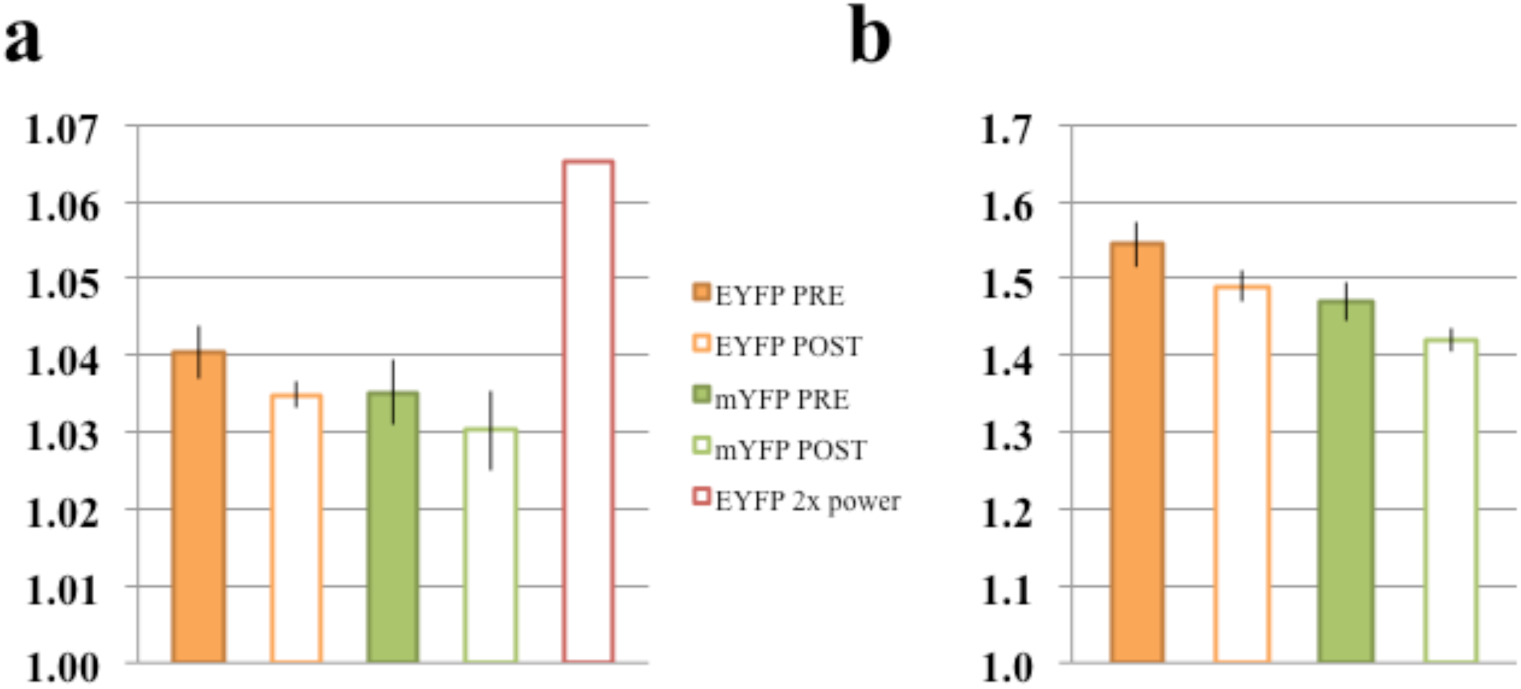
Comparison of the brightness of EYFP and mEYFP. a) Temporal brighntess analysis of EYFP and mYFP diffusing in the cytosol of a living HEK293AD cell. Note the lower EYFP brightness compared to the 2D diffusing EYFP-tagged receptors. Upon photobleaching the brighntess of both EYFP and mYFP decreases slightly, but of comparable amounts suggesting that EYFP has no more propensity to oligomerise than mYFP, which in turn may display a slightly lower intrinsic brighntess. The brightness of EYFP excited with twice the laser power is displayed as an internal control. b) Same measurements as in a performed using Spatial Brightness. Note that the brightness obtained for EYFP is now also a fraction of that obtained when looking at membrane receptors.

**SI Figure 3.**
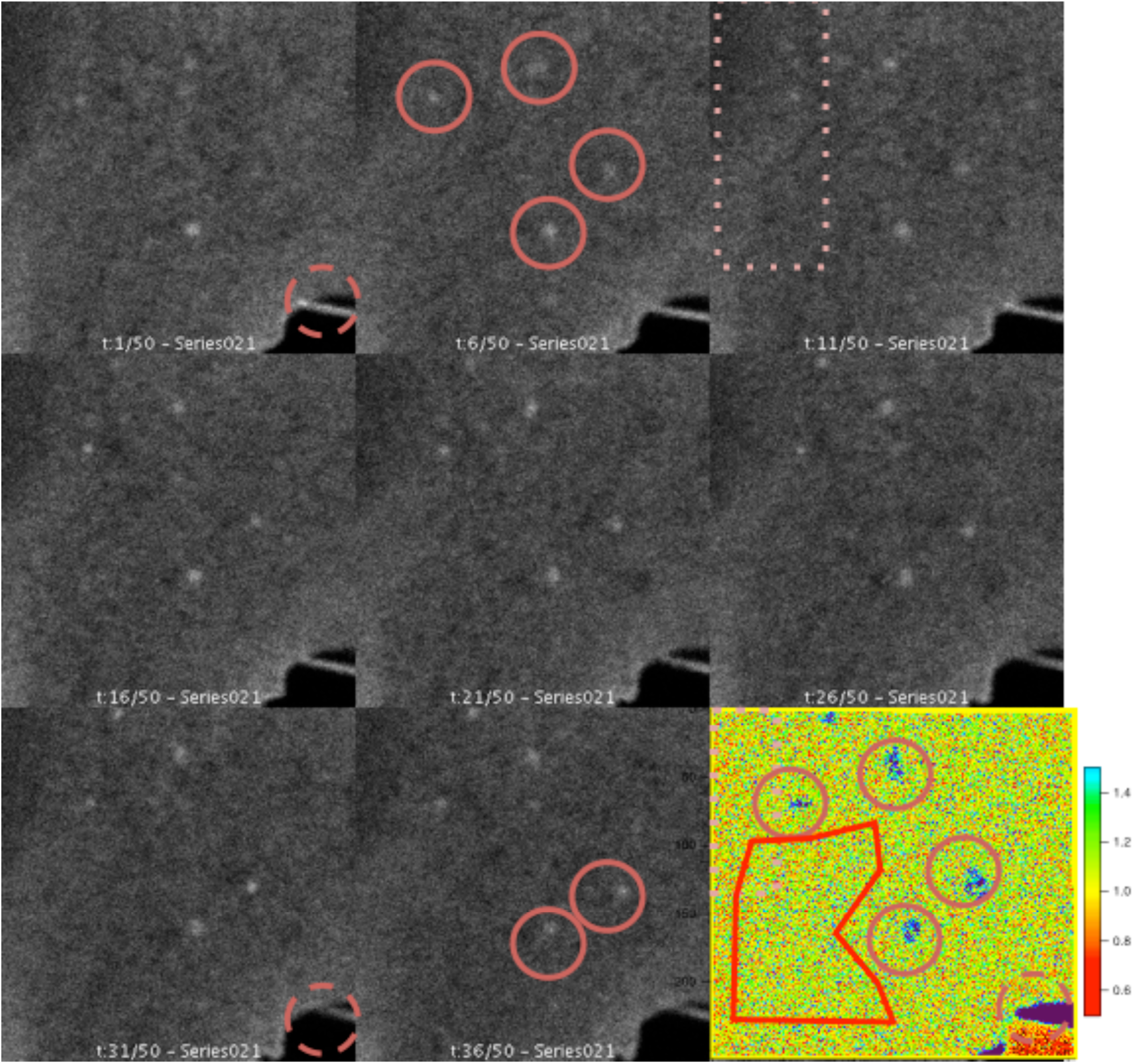
Frames obtained from a sequential imaging of the basal membrane of a Hek293 cell expressing β1-EYFP. The last frame displays the calculated apparent brightness for each pixel of the image. The decrease in intensity after 50 frames is minimal, routinely less that 10%. The dashed circle highlights a region at the edge of the cell generating high brightness due to the macroscopic movement of a protruding cell filament. Solid circles highlight four large aggregates/vescicles, giving rise to large brightness values due to their high concentration of fluorophores and small local motion. The dotted rectangle highlights a portion of the basal membrane displaying a lower intensity, but a comparable brightness with respect to the rest of the membrane. The polygon represent an example of area selection for Brightness calculation. Pixel size=50 nm, line scan rate= 200 Hz, 514 nm laser power=2%.

**SI Figure 4.**
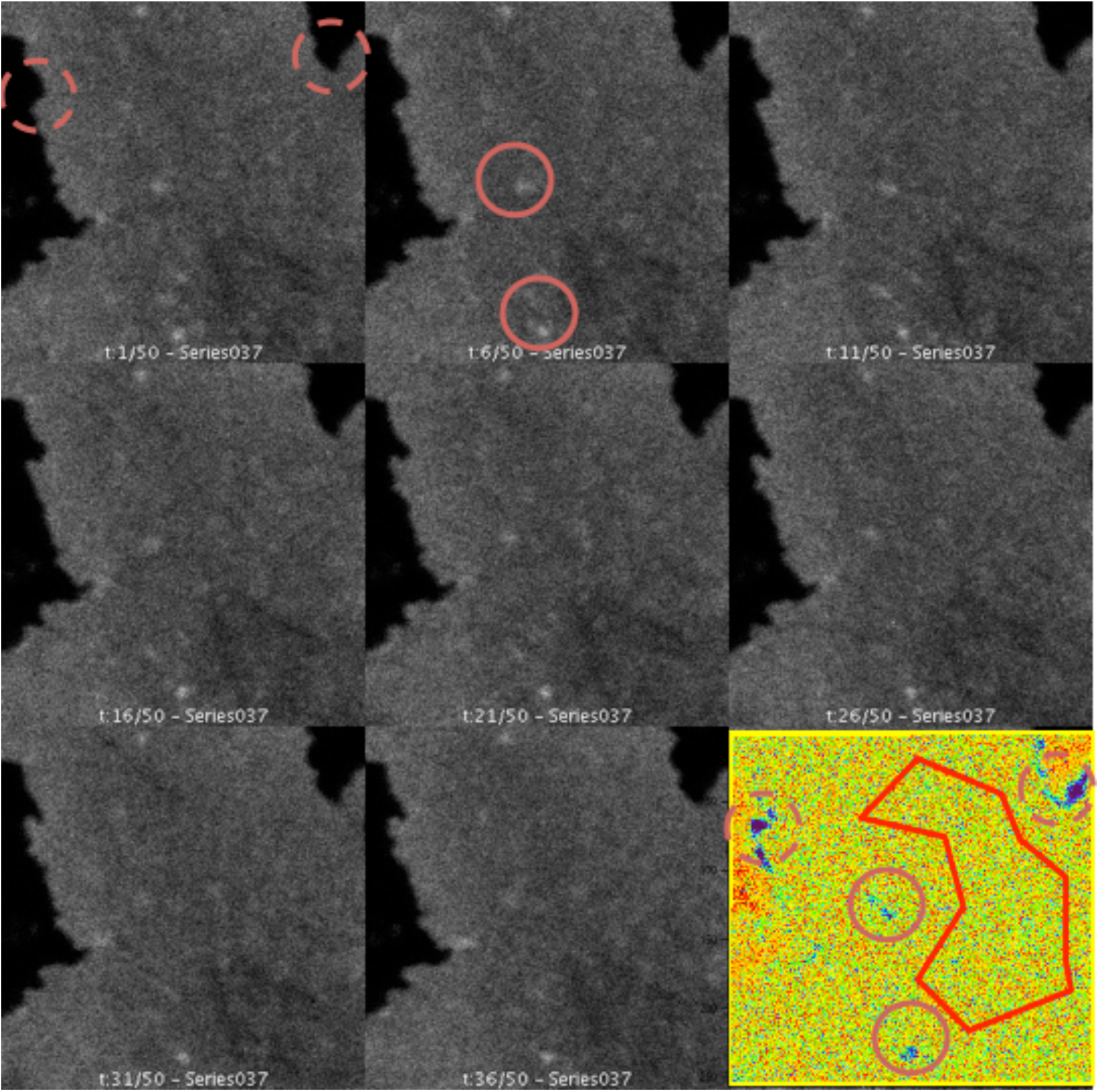
Frames obtained from a sequential imaging of the basal membrane of a Hek293 cell expressing SNAP-β1. The last frame displays the calculated apparent brightness for each pixel of the image. The intensity of the signal is virtually constant throughout the entire stack. The dashed circles highlight two regions at the edge of the cell generating high brightness due to the macroscopic fluctuation of the membrane edge. Solid circles highlight two large aggregates/vescicles, giving rise to large brightness values due to their high concentration of fluorophores and small local motion. The polygon represents an example of area selection for Brightness calculation. Pixel size=50 nm, line scan rate= 200 Hz, 488nm laser power=2.5%.

**SI Figure 5.**
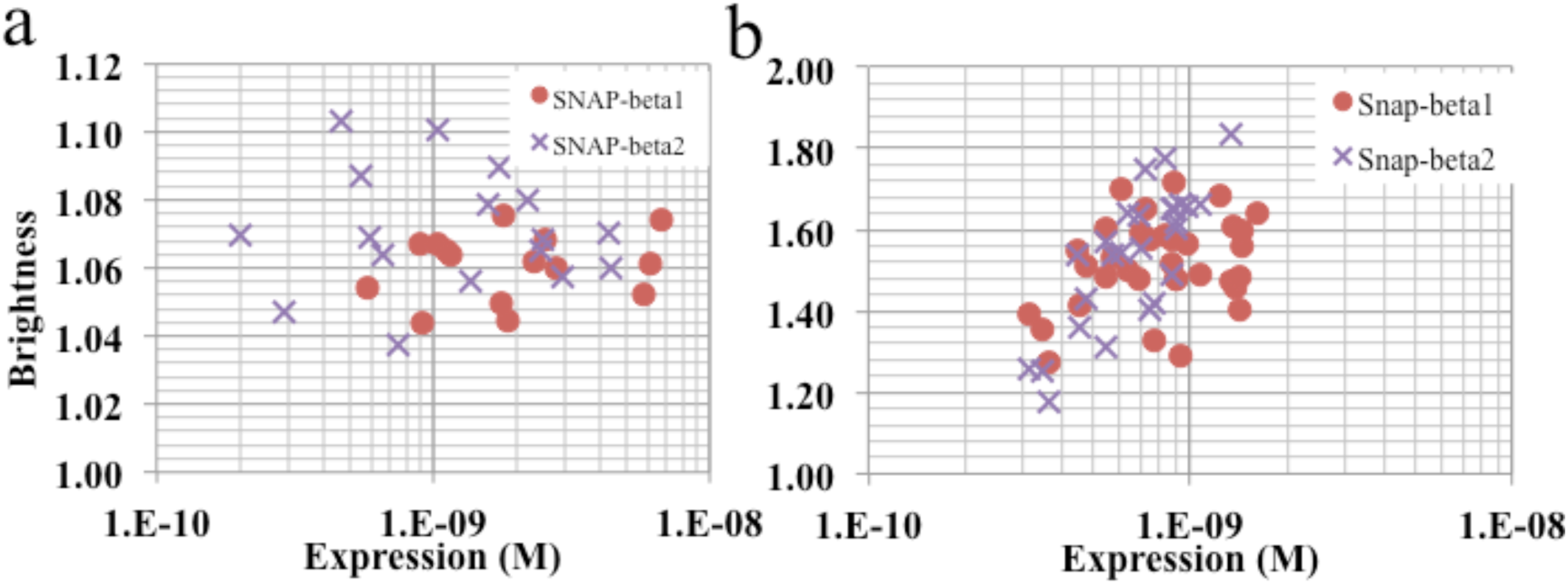
Brightness vs expression (M) for SNAP-tag labeled adrenergic receptors. a) Temporal brighntess analysis of SNAP-β1 and SNAP-β2. b) Spatial brightness analysis of SNAP-β1 and SNAP-β2.

**SI Figure 6.**
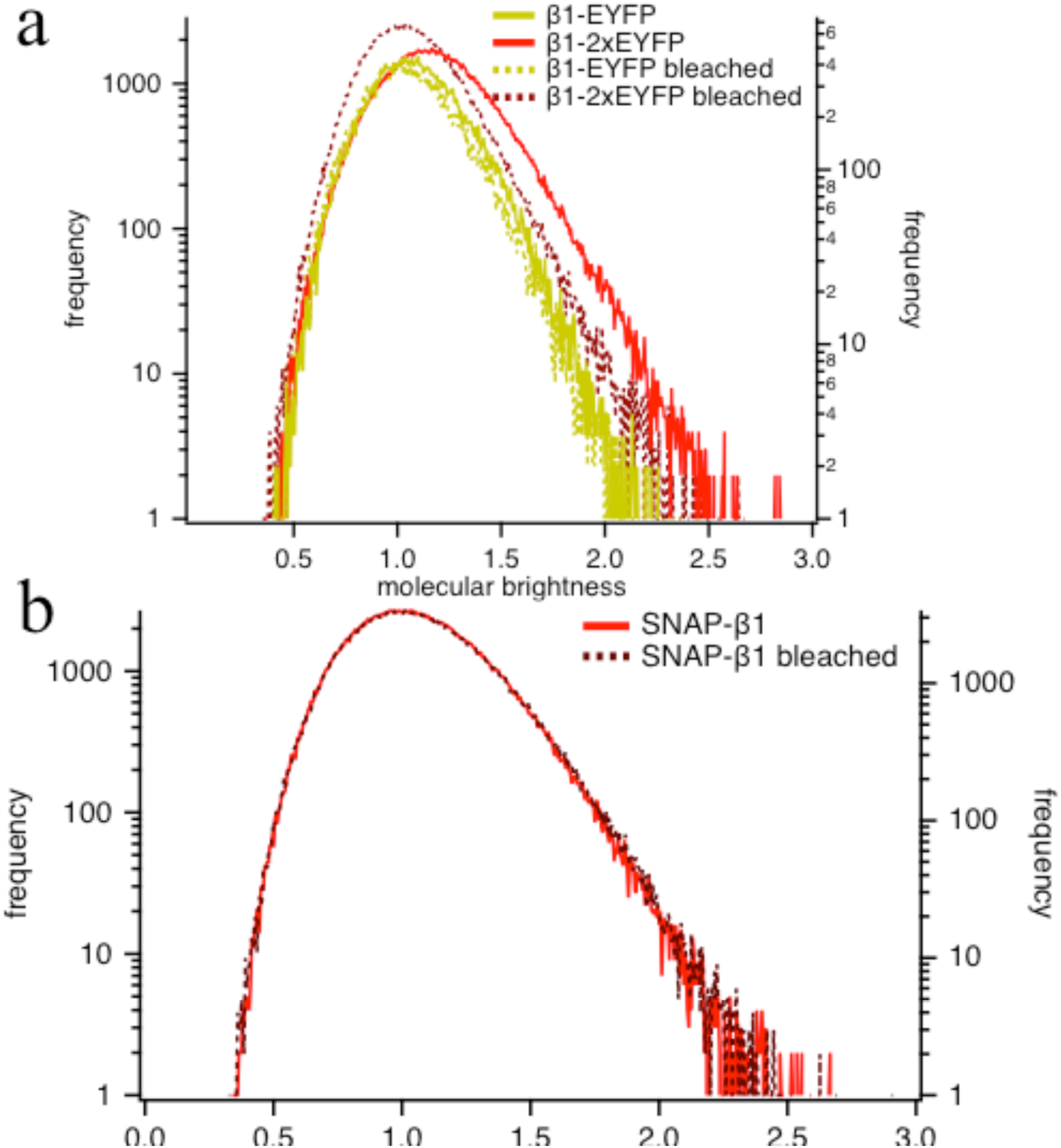
Combined Brightness histograms for three of the constructs used in this work β1-EYFP, β1-2xEYFP and SNAP-β1. a) Brightness histograms of β1-EYFP, β1-2xEYFP (_) before and after (…) the application of a whole-cell bleaching step reducing of about 50% the intensity. Post bleaching brightness for β1-EYFP is only slightly reduced, whereas a significant reduction is observed for β1-2xEYFP, shifting its histogram to the values of β1-EYFP. b) The histogram for SNAP-tag labeled β1 is virtually unaffected by photobleaching, confirming the monomeric nature of this construct. Each histogram is calculated using the brightness values from at least 10 cells. The frequency refers to pixel-occurrence of a certain brightness value.

**SI Figure 7.**
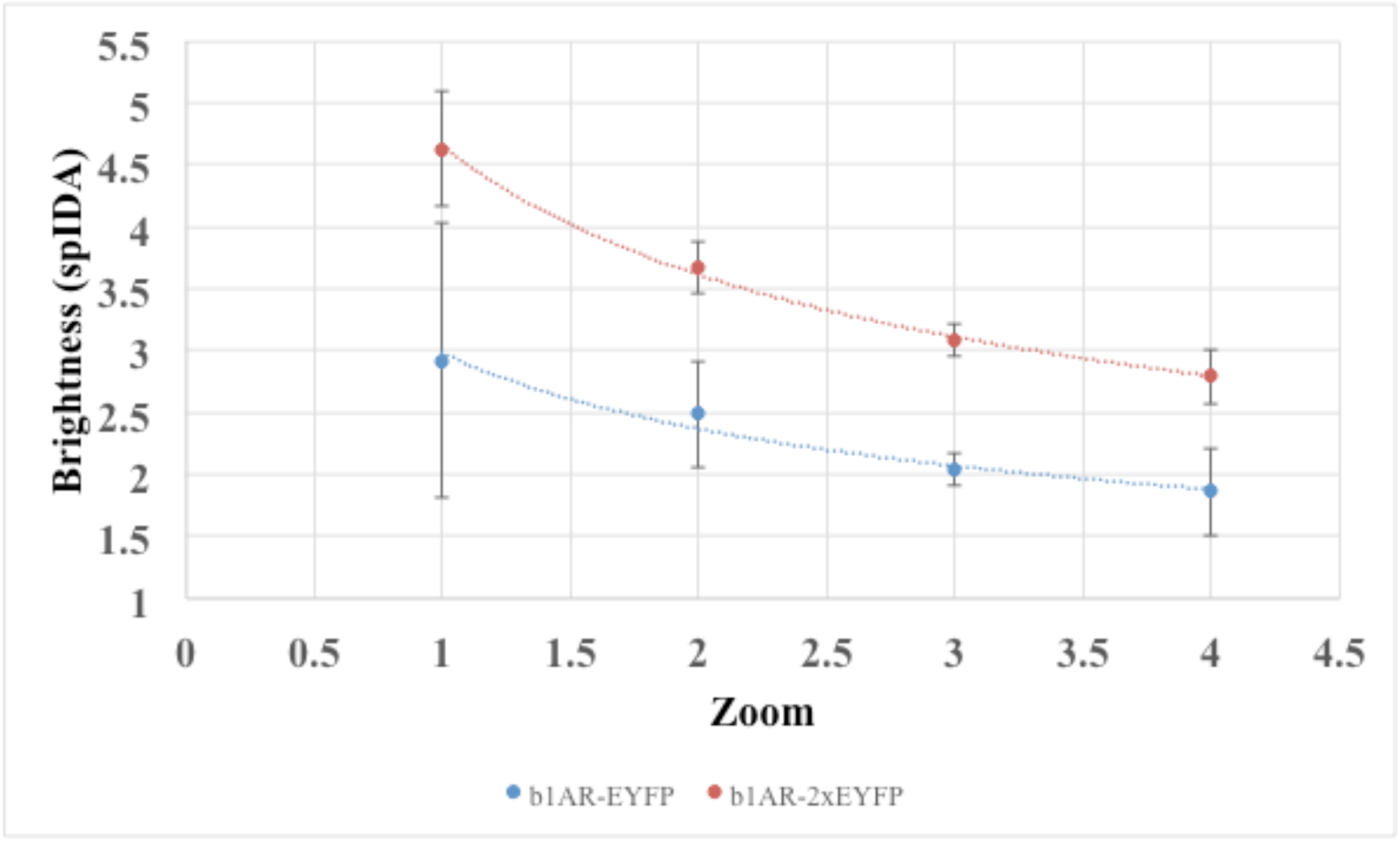
Variation of spatial brighntess as a function of the surface area of the cell membrane used for the analysis. The construct investigated is β1-EYFP. At larger areas, the brightness appears larger, and its error is also increased, since small aggregates and vesicles are included in the analysis. The smaller area selection allows an increasingly precise determination of the membrane leaflets that display homogeneous expression of the receptor.

**SI Figure 8.**
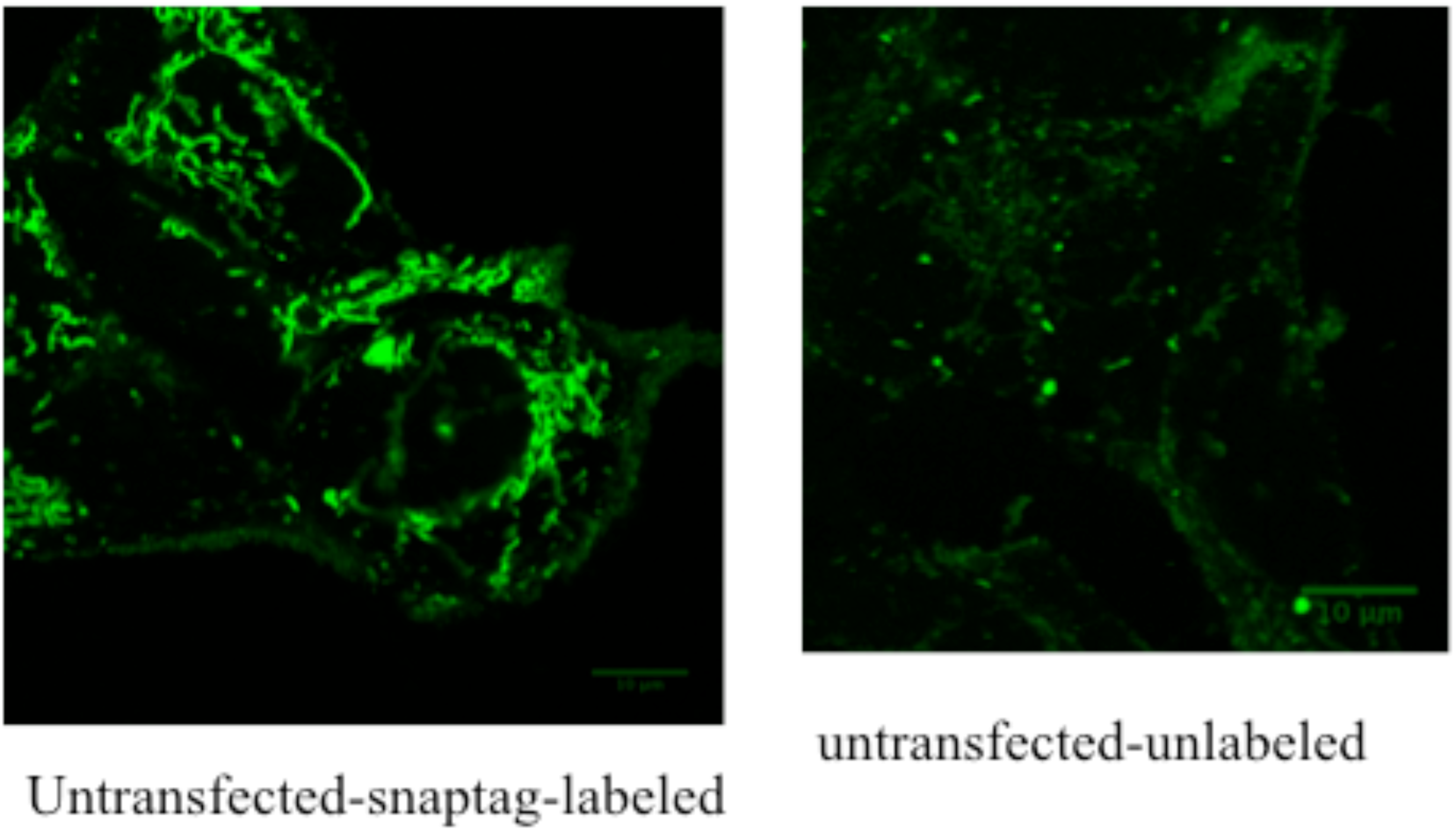
Untransfected HEK293AD cells labeled with SNAP-surface 488 according to the protocol described in the Materials and Methods. Incorporation of the SNAP-dye within cytosolic compartments can be observed in the left panel.

**SI Figure 9.**
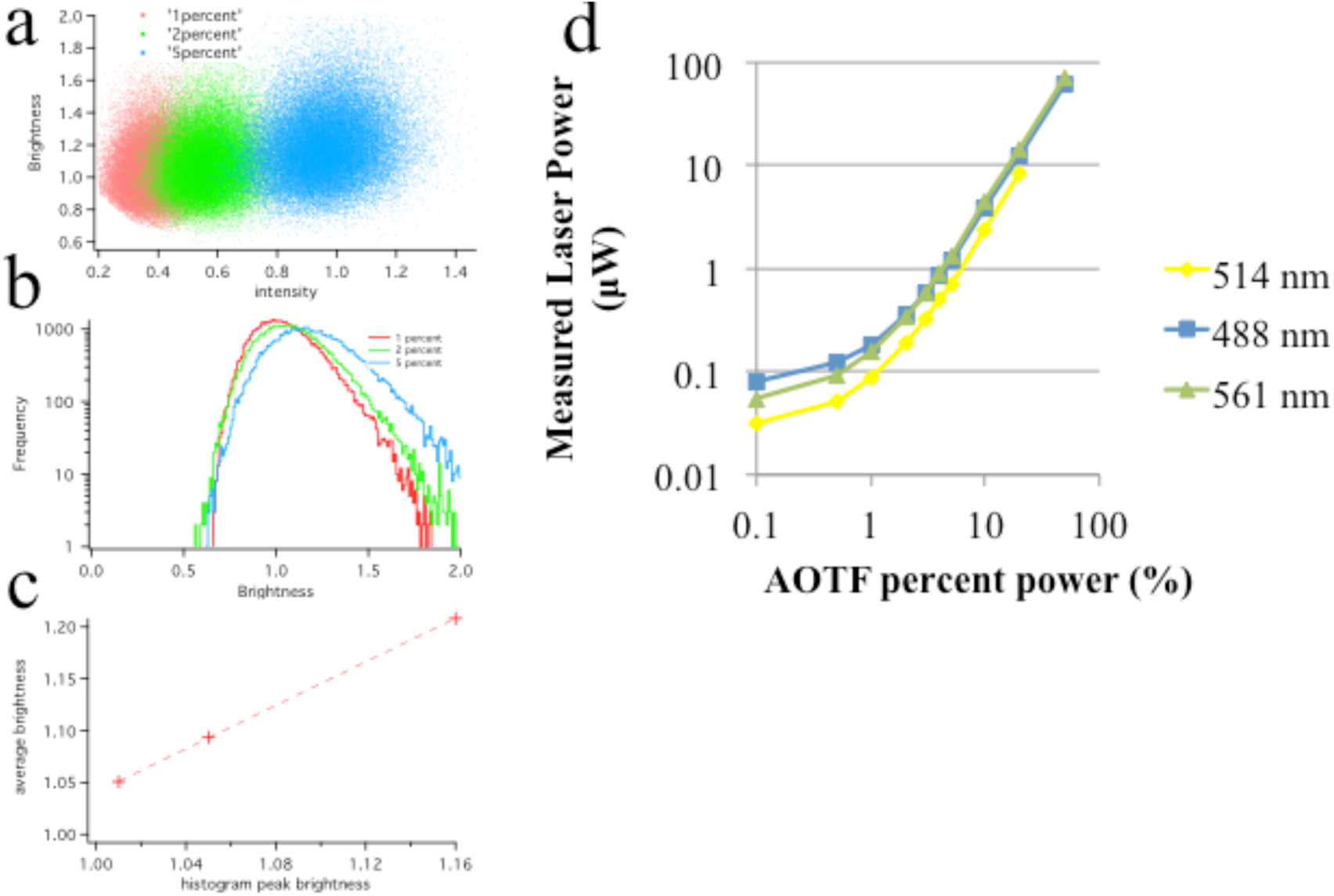
Temporal Brightness Analysis of 10 nM Alexa488 solution in a 90% Glycerol: 10% water mixture. a) Brightness vs intensity plot measured for three different laser power (determining three distinct apparent brightness values). Data were acquired at a line scan rate of 40 Hz, 256×256 pixels. b) Histogram of the brightness values displayed in a for each of the three laser power. c) correlation plot of the brightness measured averaging all the brightness points vs the brightness value obtained from the peak value in the histogram. Given the linear correlation, we used the average apparent brightness throughout the ms. d) Calibration curve of the power measured after the objective (Leica 40x, 1,25 NA, W) using a power meter (S175C Thorlabs) for three laser lines, 488 nm, 514 nm and 561 nm.

**SI Table 1.** Comparison of diffusion coefficients for c-terminally EYFP labeled receptors and n-terminally SNAP-tag labeled receptors. Diffusion coefficients were obtained from three experiments according to the iMSD methods [Di Rienzo et al. 2013]

| | $\beta 1$ - EYFP | SNAP - $\beta 1$ |
| --- | --- | --- |
| Diffusion Coefficient (0-100 ms) $\mu\text{m}^2/\text{s}$ | $0.116 \pm 0.002$ | $0.089 \pm 0.004$ |

